# AEon: A global genetic ancestry estimation tool

**DOI:** 10.1101/2024.06.18.599246

**Authors:** Naomi M Warren, Mark Pinese

## Abstract

**Background:** Genetic ancestry is a major confounding factor in genetic association studies, and it is essential to estimate and account for ancestry in genomic research. Despite this importance, accurate ancestry information is difficult to obtain, and existing methods to estimate ancestry are not designed for modern sequencing data. This methodological gap hinders the integration of genetic ancestry information in modern research, and limits progress in finding the genetic determinants of disease in diverse populations. To address this gap we present AEon, a probabilistic model-based global ancestry estimation tool, ready for use on modern genomic data.

**Results:** AEon predicts an individual’s fractional ancestral population membership, accounting for possible admixture. Unlike previous global ancestry estimation tools, AEon takes input directly from a VCF/BCF, includes default training data based on the 26 reference populations of the 1000 Genomes Project, and produces visualisation aids and diagnostics to complement data output.

**Conclusions:** AEon’s turnkey design significantly reduces the time taken to estimate ancestry from VCFs without compromising accuracy compared to ADMIXTURE analysis, with a 4.8-fold decrease in runtime for a file with 10,000 sites and 100 individuals. AEon is available at github.com/GenomicRisk/aeon.

## Background

Genetic ancestry is a major confounding factor in genetic association studies. Current association studies are heavily biased toward European populations [1,2], limiting the application of study findings [1,2,3]. Variation in linkage disequilibrium structure between populations can disrupt linkage between causal and marker alleles [3,4], and polygenic or indirect biomarkers can be confounded by uncharacterised modifiers with enriched allele frequency in specific population groups [3,5,6]. It is therefore essential to estimate and account for genetic ancestry if we hope to translate genomic research into equitable clinical utility for ancestrally diverse populations.

Despite its critical importance for equitable genomic medicine, ancestry information is not routinely incorporated in genomic analyses. This gap exists primarily because accurate ancestry data is difficult to obtain. Self-reported ancestry is not always available, and is often conflated with ethno-cultural identity, making it an inconsistent indicator of genetic background [7]. Although ancestry can be estimated directly from an individual’s genotype, and is independent of ethnic identity, such genotype-derived ancestry can be difficult to estimate in practice.

Computational methods have been developed to estimate genetic ancestry from genotype data, but current tools suffer from significant practical limitations that hinder their adoption in modern genomic workflows. Most existing tools either do not use the modern defacto VCF input format (ADMIXTURE [8], iAdmix [9]), rely on external infrastructure (gnomAD’s AE tool on Hail [10], Rye [11]), require phased genotypes as input (FLARE [12], MOSAIC [13], GNOMIX [14]), or employ difficult-to-obtain individual-level genotype data from population reference individuals (LASER [15]). The SNVstory method [16] addresses several of these limitations, but can only identify the major ancestral population in an individual, making it unsuitable for admixed individuals – an increasingly common scenario in modern clinical cohorts.

Genetic ancestry inference methods can be broadly categorized along two dimensions: global versus local approaches, and supervised versus unsupervised methods. Global methods model the entire genome as a homogeneous mixture of ancestral populations, while local methods identify the ancestral origin of specific genomic segments. Supervised methods estimate an individual’s ancestry as a mixture of predefined reference populations, whereas unsupervised approaches infer ancestral populations directly from the data. For clinical genomics applications, global supervised approaches are ideal: they work efficiently on unphased data (unlike local methods) and provide interpretable, consistent ancestry estimates based on established reference populations (unlike unsupervised approaches). Unfortunately, no existing tool provides user-friendly global supervised ancestry estimation for modern sequencing data formats.

To address this gap, we have developed AEon, a probabilistic model-based tool for supervised global ancestry estimation, ready for use on modern genomic data. AEon predicts fractional population membership of input samples given allele frequency data from known populations, accounting for possible admixture. It is simple to install and run, implemented in Python, and utilises existing mature libraries to process input and perform statistical modelling. AEon takes input directly from a VCF/BCF and outputs estimates in CSV format, enabling easy integration with existing software and pipelines. The tool also includes a default population reference file to provide turnkey ancestry estimation with no further configuration. However, the option to modify or replace this reference file is available should users wish to tailor the tool for their specific input data and questions of interest.

AEon removes the barriers preventing genetic ancestry estimation from being widely integrated into genome analysis pipelines, thereby enabling and accelerating ancestry-informed genetic research. Here we present a high-level overview of AEon, describe its performance and application to modern genomic sequencing data relative to the traditional ADMIXTURE tool, and make the tool and its databases available for use.

## Implementation

### Approach

AEon estimates fractional population membership for individuals based on their genotype calls and a population-specific allele frequency reference.

Briefly, AEon employs `pysam` to read input genotypes in VCF/BCF format, and converts genotypes to dosages of pre-specified ancestry-informative alleles. These dosages are then fit to a Gaussian mixture model by maximum likelihood using `pyro` [17], to yield a per-individual population mixture estimate. This fit requires a reference table giving frequencies of each allele in each population of interest. We supply a default table for GRCh38 across the 1000 Genomes (G1K) Project populations, but AEon also supports user-supplied tables.

As output AEon produces per-individual population mixture estimates in CSV format.

### Statistical Model

Following the ADMIXTURE approach [8], AEon models an individual as a mixture of prespecified ancestral populations. Specifically, denote the count of non-reference alleles of an individual at biallelic locus *j* as *G*_*j*_ ∈ {0, 1, 2}. Under the gene pool model, we suppose each *G*_*j*_ is drawn independently from a categorical distribution with probabilities 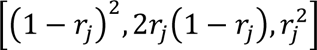 for *G*_*j*_ = [0, 1, 2], where *r*_*j*_ is the non-reference allele frequency for the gene pool from which the individual was sampled. We model the gene pool frequency for this individual *r*_*j*_ as a mixture of ancestral frequencies, *r*_*j*_ = ∑_*i*_ *f*_*ij*_*p*_*i*_ where *p*_*i*_ ∈ [0,1] is the fraction of ancestral population *i* contributing to this individual, and *f*_*ij*_ is the non-reference allele frequency at locus *j* in ancestral population *i*, ∑_*i*_ *p*_*i*_ = 1. Given prespecified *f*_*ij*_ and supplied individual *G*_*j*_ from genotyping, AEon estimates the full vector of population memberships *p* by maximum likelihood.

### Generating the Population Reference

AEon includes a default reference table of ancestry-informative allele frequencies from the 2,504 unrelated individuals in the G1K Project Phase 3 dataset [18], sampled from 26 populations grouped into 5 superpopulations. This table was based on a list of 133,872 alleles previously determined to be ancestry-informative markers across the 1000 Genomes populations based on Hardy-Weinberg analysis [19]. The list was pruned to remove difficult to sequence regions and closely-linked loci. We further filtered the list to remove alternate alleles absent from GRCh38 and loci missing from the 1000 Genomes call set to yield a final list of 128,097 alternate alleles. One sample from each of the 26 populations was set aside to form a test set (Table S1), and the remaining samples in each population were used to calculate allele frequencies at each locus to form the final reference table.

### Testing on 1000 Genomes populations

Ancestry estimation with AEon was tested on WGS data from the 1000 Genomes Project using the 26 samples set aside from the reference in the previous step. Genotype calls for these 26 individuals were provided to AEon as an indexed BCF using default flags. To evaluate the impact of number of loci on performance, AEon models were fit with either a random subset of 1,000, 2,000, 5,000, 10,000, 20,000, 50,000 or 100,000 loci, or all 128,907 default reference loci from the reference allele frequency file. The estimated ancestry fractions provided as output from these tests were then compared against the labelled superpopulation assigned by the G1K database for each sample.

### Testing on Simulated Admixed Individuals

Since all individuals from the G1K database are intended to be representative of specific populations, we simulated admixture to test AEon’s performance on individuals with genetic ancestry from multiple population groups at different proportions. First, mixtures of CEPH European Ancestry in Utah (CEU) and Sri Lankan Tamil in the UK (STU) were generated by treating all loci as unlinked and randomly passing on one allele from each parent to the simulated child. We chose this independent sampling strategy as loci used by AEon are sparsely sampled to ensure they are likely unlinked. The simulated CEU50:STU50 individual was then designated as a new ‘parent’ along with a Han Chinese South (CHS) individual to generate 20 samples with CEU25:STU25:CHS50 mixed ancestry. The same number of simulated individuals were produced with a mix of CEU12.5:STU12.5:CHS75 and CEU37.5:STU37.5:CHS25 ancestry by combining the CEU25:STU25:CHS50 sample with either a CEU50:STU50 or CHS100 sample respectively (Figure 1b). After preliminary testing we found that estimated ancestry fractions were stable using at least 10,000 loci as markers; therefore these simulated individuals were run through AEon using eight randomly sampled subsets of 10,000 loci, and average predictions were calculated. Given that these individuals did not belong to a single population group, their estimated ancestry fractions from each trial were compared against the G1K labelled fractions using L2-distance between ancestry fraction vectors.

**Figure 1.**
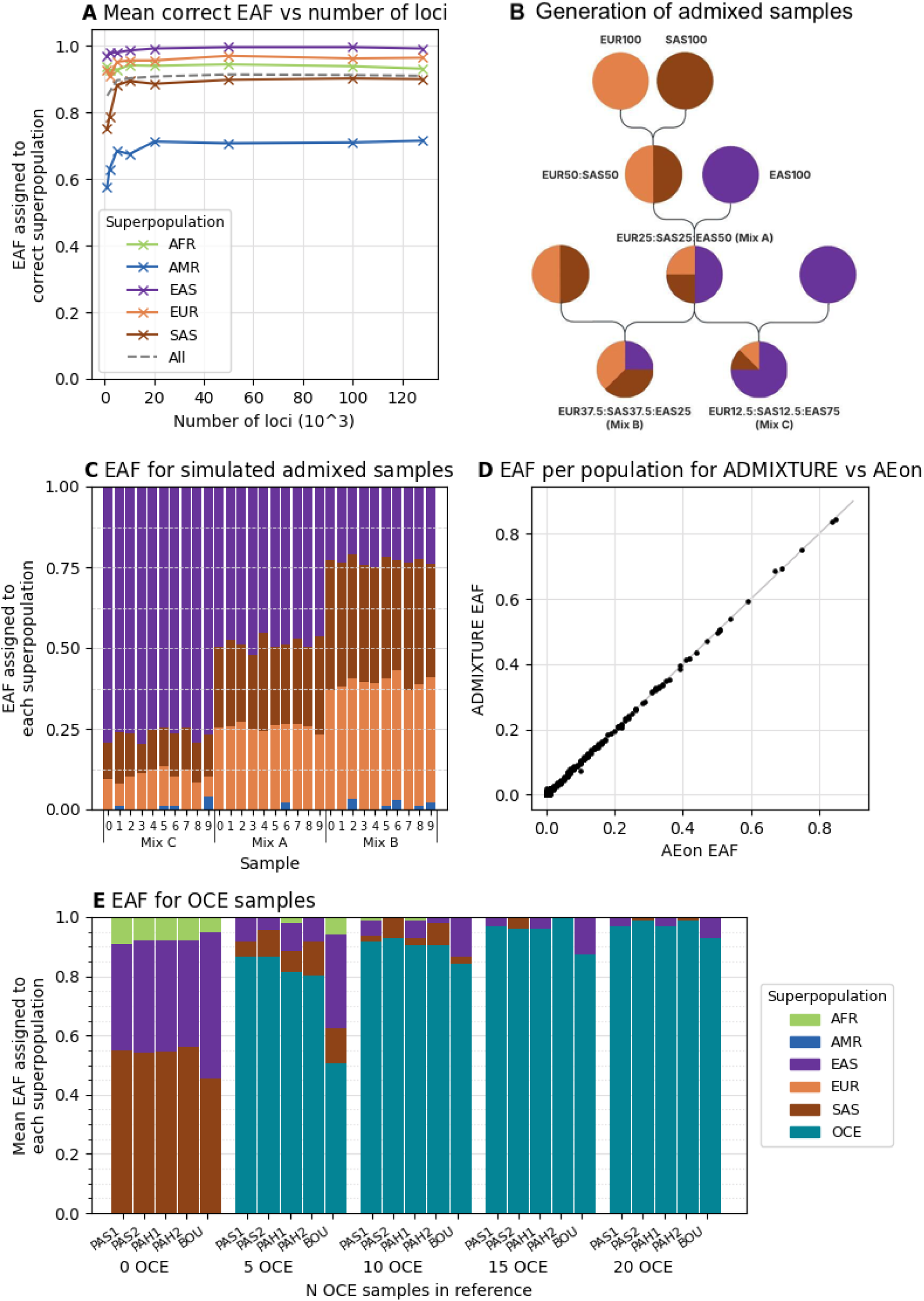
High accuracy of superpopulation composition can be achieved with sufficient loci, with estimates reaching optimal accuracy from approximately 10,000 loci **(a)**. AEon predicts suitable superpopulation ancestry fractions for admixed individuals **(c)**, the lineage of which is shown in **(b)**. EAFs are nearly identical between ADMIXTURE and AEon, lying along the line y=x. Each point represents the EAF assigned to a population for a given sample, with 26 points per sample **(d)**. AEon is able to incorporate new population information, allowing samples from previously underrepresented populations to be characterised **(e)**. Note that the colour legend for superpopulations is consistent across all plots in this figure.

### Comparison to ADMIXTURE

Having validated AEon’s results against the labelled populations, we also compared its performance to the existing ancestry estimation tool ADMIXTURE, used in projection mode (ADMIXTURE’s supervised inference mode, selected with the -P command line flag).

First ADMIXTURE required an initial run on the reference set using `--supervised` mode to generate a ‘.P’ population reference file. To run ADMIXTURE on the test samples, a pre-processing script that calls `bcftools` and `plink` was created to convert the initial BCF into the required format (see Additional file 1), after which ancestry estimation for the test samples was performed using ADMIXTURE’s projection mode with the ‘.P’ file generated earlier from the reference set. Maximum memory usage, maximum CPU, and real time spent were recorded for both the pre-processing step and the predictive projection-mode ADMIXTURE run using the same selection of loci (between 1,000 and 100,000) as the AEon trials. Tests were also conducted varying the number of samples in the input file, all with 10,000 loci. We compared these resource metrics between tools, as well as the per-sample population mixtures predicted by each tool, on a MacBook Pro machine (Apple M2 Pro chip with 10-core CPU (6 performance cores, 4 efficiency cores), 16GB memory). Note that the time and memory required for the initial supervised run was not included in these comparisons.

### Extending to other reference populations

To establish AEon’s performance when extending to new reference populations, we tested its performance on Oceanian samples from the Human Genome Diversity Project [20] (HGDP) and Simons Genome Diversity Project [21] (SGDP). The genotypes of 7 Papuan Sepik (PAS), 7 Papuan Highlands (PAH) and 11 Bougainville (BOU) individuals were downloaded in VCF format. 2 PAS, 2 PAH and 1 BOU individuals were reserved as test samples. The remaining 5 PAS, 5 PAH and 10 BOU individuals were used to generate allele frequencies per population using `bcftools`. To understand how reference sample size impacted the quality of results, four new reference files were generated with increasing Oceanian input. The ‘5 OCE’ reference was created by taking the default AEon G1K reference file and adding a new ‘PAS’ column, which contained the allele frequencies calculated from the 5 reference PAS samples at AEon’s 128,097 loci. The ‘10 OCE’ reference took this new file and added an additional ‘BOU’ column, where the allele frequencies for the BOU population were calculated using only 5 of the reference BOU samples. PAH allele frequencies from 5 individuals were added in a new column to produce the ‘15 OCE’ reference. Finally, the ‘20 OCE’ reference was created by replacing the BOU column with allele frequences calculated using all 10 reference BOU samples. The content of these allele frequency references, testing the effect of including different numbers of Oceanian individuals in the reference table, is summarised in Supplementary Table S2. The addition of new populations required adding corresponding rows to the tabular ‘population labels’ file, which contains the superpopulation assignment for each population. For each new reference file, AEon was run on the 5 test samples at 10,000 loci, using the flags *`-a new_reference_afs.txt -- population_labels G1K_OCE_labels.txt`*.

## Results

Existing global genetic ancestry estimation tools require significant hands-on time to apply on modern sequencing data, limiting their uptake. AEon streamlines global ancestry estimation by running directly from VCF files. This decreased complexity reduces analysis burden and eliminates error-prone data conversion steps. Here we demonstrate that this simplicity does not come at the expense of accuracy, and that AEon yields accurate ancestry estimates in single-population and admixed samples, competitive with gold-standard global approaches such as ADMXITURE.

### AEon identifies the most significant superpopulation ancestry in an individual

We first evaluated AEon’s ability to correctly infer the largest contributing ancestry in an individual. We applied AEon to 26 held-out samples from the 1000 Genomes Project, and measured the concordance between AEon’s largest estimated ancestry fraction (EAF) and the sample’s labelled superpopulation.

AEon correctly identified the major contributing superpopulation in 25 out of 26 cases, independent of the number of ancestry-informative loci supplied (Figure 1a). The remaining sample, labelled in 1000 Genomes as Colombian (CLM), was assigned primarily European superpopulation ancestry by Aeon, followed by the American superpopulation. While as few as 1,000 loci could be used to assign the largest proportion of ancestry to the correct superpopulation for most cases, this proportion and hence confidence in the prediction increased when more loci were provided, with diminishing returns beyond 10,000 loci (Figure 1a).

While the CLM sample was primarily assigned European ancestry at the superpopulation level, its dominant ancestral component at the population level was CLM when 20,000 or more loci were used for inference (Figure S1). AEon’s prediction of a large fraction of European ancestry in the CLM individual was consistent with the known admixed ancestral composition in this group [22]. Given the potential for AEon to detect and quantify levels of admixture, we then explicitly investigated AEon’s ability to accurately estimate EAF in individuals with defined admixture.

### AEon accurately estimates ancestry of admixed individuals

To evaluate AEon’s ability to estimate known admixture, we simulated offspring from random pairings of individuals from the 1000 Genomes CEU, STU and CHS populations and their offspring (figure 1b). This procedure generated offspring with three different admixture patterns: 25% CEU + 25% STU + 50% CHS, 37.5% CEU + 37.5% STU + 25% CHS, or 12.5% CEU + 12.5% STU + 75% CHS. With only 10,000 loci as input, AEon accurately and consistently predicted the known ancestral composition of these complex admixtures (Figure 1c). These results establish AEon’s ability to estimate ancestry composition in admixed samples and supports the use of 10,000 loci for this application. To further characterise AEon’s performance, we compared its estimates and computational performance to the gold-standard ADMIXTURE method.

### AEon matches ADMIXTURE’s accuracy, with enhanced performance

AEon ancestry estimates were equivalent to those from ADMIXTURE. Across the 26 held-out test samples from the 1000 Genomes Project, AEon and ADMIXTURE assigned nearly identical ancestry fractions to each population, with a mean difference of 0.00 and maximum difference of 0.027 for a single population when considering all 128,907 loci (Figure 1d).

Although AEon demonstrated equivalent accuracy to ADMIXTURE, it was more computationally efficient. AEon ancestry estimation was significantly faster than ADMIXTURE in wallclock time, both as a function of number of loci and the number of samples (Figure 2a,b). This was largely due to the costly format conversion step required for ADMIXTURE (Table S3), as well as AEon’s more efficient utilisation of computational resources (Figures 2c,S2). The initial supervised ADMIXTURE run that was required to generate a reference file presented an additional one-off time cost, and was not included in these comparisons. Although AEon did require more system memory for its computation, it had superior memory scaling to ADMIXTURE, and the RAM requirements for either tool are unlikely to be limiting on modern hardware (Figure 2d). As AEon operates on samples serially, its RAM usage is effectively independent of cohort size.

**Figure 2.**
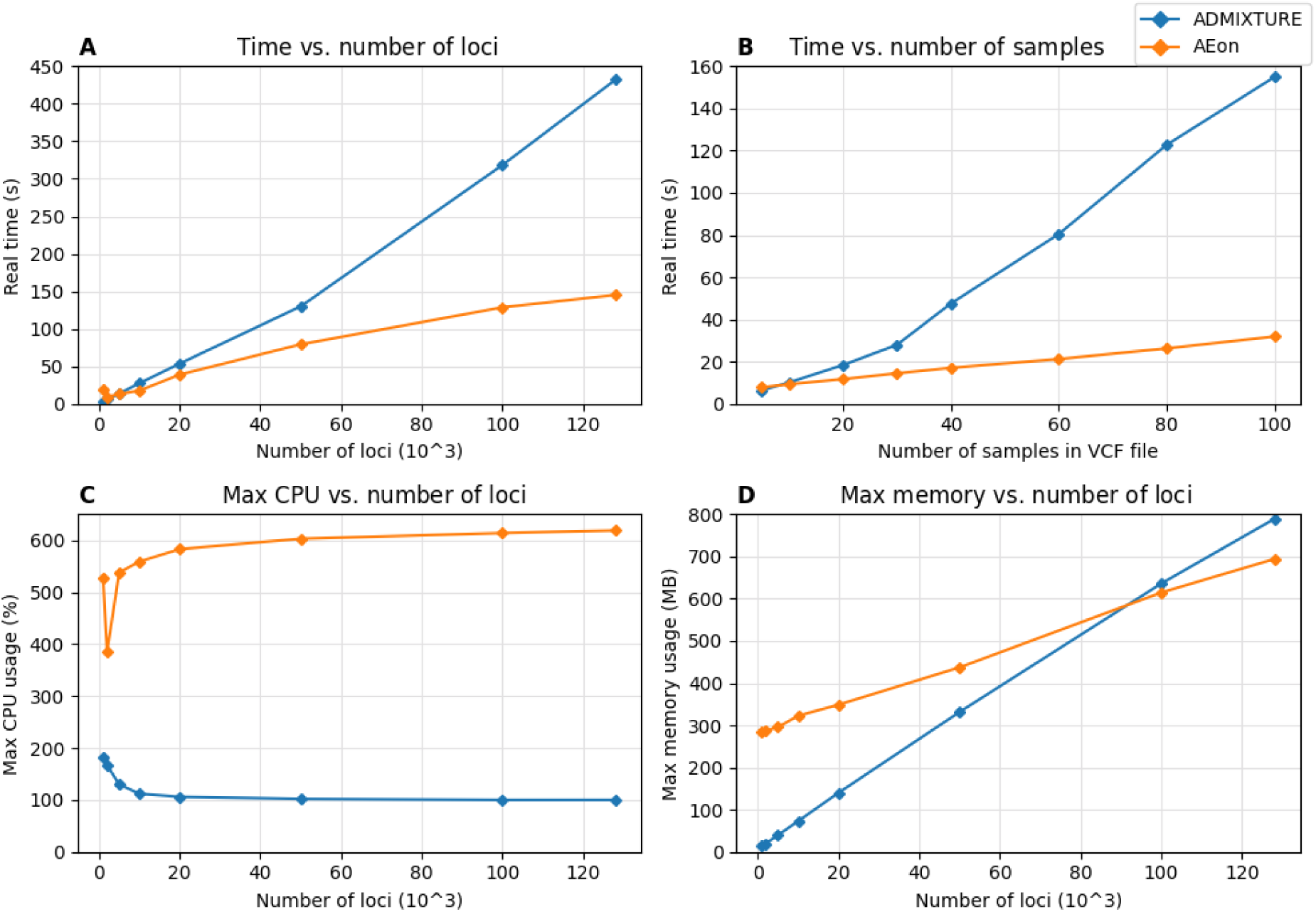
Comparison between the resources used by AEon (orange) and ADMIXTURE (blue). AEon is faster than ADMIXTURE, and its runtime scales better with increasing locus count **(a)** or sample count **(b)**. AEon automatically makes use of available CPU regardless of loci count **(c)**. AEon’s memory usage also scales better with loci count **(d)**.

In summary, AEon simplifies the process of estimating ancestry, decreasing the time required to perform analysis both in terms of wallclock time and analyst effort (Figure S3). AEon’s simplicity does not compromise accuracy, resulting in ancestry estimates that are of the same quality as the current standard tool ADMIXTURE.

### AEon readily extends to new populations with few reference individuals

AEon’s underlying model relies on the data provided in its reference file, allowing the user to interpret results within the context of predefined populations. However, this requires that AEon’s reference file contains information for all populations of interest. AEon is designed to be extensible to accommodate new reference populations, and to demonstrate this capability and its data requirements, we undertook a study examining AEon’s ability to quantify Oceanian ancestry.

As a baseline, we first evaluated AEon’s behaviour when supplied test individuals of Oceanian ancestry, a population group that is absent from AEon’s default G1K reference population file. As expected, applying AEon to Oceanian individuals using the default G1K reference file led to inaccurate results, with Oceanian individuals assigned a mixture of South Asian, East Asian, and African ancestry (Figure 1e, left bar group).

We then evaluated how rapidly AEon’s ancestry estimates became accurate when it is supplied increasing numbers of Oceanian individuals in its reference file. We generated reference files that included varying numbers of individuals of Papuan Sepik (PAS), Papuan Highlands (PAH), and Bougainville (BOU) ancestry, and tested AEon’s performance on held-out Oceanian samples. Inclusion of reference allele frequencies generated from only 5 PAS samples resulted in an estimated Oceanian ancestry fraction of over 0.85 for the test PAS samples, and a fraction above 0.8 and 0.5 for the PAH samples and BOU sample respectively (figure 1e, 2^nd^ bar group). This indicates that AEon is capable of accurately identifying underrepresented ancestral populations with a minimal amount of reference data. Including reference BOU allele frequencies significantly improved the fraction of Oceanian ancestry assigned to the BOU sample and marginally improved the assigned Oceanian fraction for the PAS and PAH samples (Figure 1e, 3^rd^ bar group). Further improvement was observed with the addition of more reference samples (Figure 1e, right two bar groups). We confirmed that the inclusion of Oceanian populations in the reference file had minimal impact on the assignment of ancestry fractions for G1K test samples (Figure S4).

This demonstrates that AEon can accurately identify population membership even when given reference frequencies based on as few as five individuals, with negligible effect on other populations. Importantly, inclusion of allele frequencies from other populations within the same superpopulation also improved correct superpopulation assignment (Figure 1e, 2^nd^ panel), indicating that even if a perfectly matched population reference is not available, a reference based on an ancestrally similar group can be used to generate accurate ancestry estimates using AEon.

## Discussion

We have presented AEon, a tool to estimate global genetic ancestry from modern sequencing data. AEon has been engineered for ease-of-use and flexibility, minimising data handling overhead and the potential for errors. As an updated implementation of the ADMIXTURE global ancestry model, AEon complements recent methods for ancestry classification, and addresses the usability gap left by current ancestry mixture modelling tools.

AEon’s primary contribution is as an ancestry mixture modelling method that works natively on modern genotype data formats. Classical methods for this modelling, such as ADMIXTURE, were designed for genotyping microarray data. Such microarray data differ significantly from modern genotyping-by-synthesis (GBS) in both file format and interpretation, particularly with respect to the handling of ungenotyped loci and homozygous reference loci [23]. Consequently, although it is possible to apply ADMIXTURE and related methods to modern GBS data in VCF format, this comes with significant manual (Figure S3) and computational overhead (Figure 2), leading to complex and error-prone workflows. By working natively with VCF data, AEon circumvents these issues and provides a highly performant, lightweight, and convenient framework to estimate ancestry from genomic data.

A key advantage of AEon is its easy and flexible setup. AEon has straightforward single-step execution, lightweight dependencies, and comes with a default ancestry reference table. The inclusion of a reference ancestry table is a notable benefit of AEon, enabling new users to generate accurate ancestry estimates for 1000 Genomes populations with minimal effort. This simplicity of AEon does not come at the expense of flexibility, and AEon accepts custom ancestry tables if required. Notably, we have shown that AEon provides accurate ancestry estimates using as few as five individuals from a new reference population. To our knowledge, this flexibility and data efficiency is unique, and supports the use of AEon to estimate ancestry in undersampled populations. With our findings that results can be significantly influenced by reference database mismatch (e.g. AMR admixture, Figures 1a,S1) or incompleteness (figure 1e), this capability of AEon enables equitable genomics for understudied human populations.

As a modern implementation of the classical ancestry mixture model, AEon complements other new tools. For example, the SNVStory method, and the ancestry estimation module of Hail [10], also aim to ascertain ancestry from modern GBS data. However, these methods are designed as classification tools, which assign individuals to a single ancestry. For populations with low admixture this is an appropriate methodology, however it is unsuitable for characterising admixed individuals. AEon, like the classical ADMIXTURE and STRUCTURE methods, natively fits ancestral mixtures and so naturally and accurately handles admixture, which is commonly encountered in modern genomic cohorts.

While AEon represents a substantial improvement in usability and performance over current approaches, it has two notable limitations. Firstly, as a supervised method, AEon can only identify ancestry from populations represented in its reference database. Although we have demonstrated that AEon can accurately infer ancestral composition from as few as five population reference individuals, this need for a reference population is a crucial property that must be considered when applying the method in practice. Secondly, as a global ancestry estimation method, AEon cannot deduce the ancestral source of specific haplotypes observed in an individual, which may be relevant to clinical interpretation of genomic findings.

## Conclusions

AEon accurately and rapidly estimates global genetic ancestry from WGS data, determining the fractional ancestral populations contributing to an individual. Built on python to run with a single command and including default reference data, it allows meaningful information to be extracted from genotype data immediately upon installation, while also being readily extensible to new reference populations.

## Supporting information

ADMIXTURE pre-processing script

## Availability and requirements

**Project name:** AEon

**Project home page:** github.com/GenomicRisk/aeon

**Operating systems:** platform independent (can be installed from GitHub or docker images can be accessed from https://hub.docker.com/r/naomiwren/aeon)

**Programming language:** Python (tested on Python 3.10)

**Other requirements:** Installation from the GitHub repository requires pip. Use of a virtual environment (e.g. conda) is also recommended.

**Licence:** Apache-2.0 licence

## List of abbreviations

AE: ancestry estimation
CPU: central processing unit
EAF: estimated ancestry fraction
GBS: genotyping-by-synthesis
G1K Project: 1000 Genomes Project
HGDP: Human Genome Diversity Project
SGDP: Simons Genome Diversity Project

For population name abbreviations, consult Supplementary Table 1.

## Declarations

### Ethics approval/consent to participate

Not applicable.

### Consent for publication

Not applicable.

### Availability of data and materials

The 1000 Genomes data analysed during the current study was downloaded from https://ftp-trace.ncbi.nlm.nih.gov/1000genomes/ftp/1000G_2504_high_coverage/working/20201028_3202_raw_GT_with_annot/.

HGDP data was accessed from https://gnomad.broadinstitute.org/downloads#v3-hgdp-1kg.

SGDP data was accessed from the Seven Bridges Cancer Genomics Cloud at https://cgc.sbgenomics.com/u/sevenbridges/simons-genome-diversity-project-sgdp.

### Authors’ contributions

NMW wrote the software and wrote the manuscript. MP conceived the software and contributed to the manuscript. All authors read and approved the manuscript.

## Acknowledgements

Not applicable.

## Supplementary Data

### Tables

**Table S1.** Selected test samples from the 1000 Genomes Project.

| Superpopulation | Population Code | Population Name | Sample ID |
| --- | --- | --- | --- |
| **African** | ACB | African Caribbean in Barbados | HG01879 |
|  | ASW | African Ancestry in South West USA | NA19625 |
|  | ESN | Esan in Nigeria | HG02922 |
|  | GWD | Gambian in Western Division - Mandinka | HG02461 |
|  | LWK | Luhya in Webuye, Kenya | NA19017 |
|  | MSL | Mende in Sierra Leone | HG03052 |
|  | YRI | Yoruba in Ibadan, Nigeria | NA18486 |
| **American** | CLM | Colombian in Medellín, Colombia | HG01112 |
|  | MXL | Mexican Ancestry in Los Angeles<br>CA, USA | NA19661 |
|  | PEL | Peruvian in Lima, Peru | HG01565 |
|  | PUR | Puerto Rican in Puerto Rico | HG00551 |
| **East Asian** | CDX | Chinese Dai in Xishuangbanna,<br>China | HG00759 |
|  | CHB | Han Chinese in Beijing, China | NA18525 |
|  | CHS | Han Chinese South | HG00403 |
|  | JPT | Japanese in Tokyo, Japan | NA18939 |
|  | KHV | Kinh in Ho Chi Minh City, Vietnam | HG01595 |
| **European** | CEU | CEPH collection - Utah Residents<br>with Northern and Western<br>European Ancestry | NA06984 |
|  | FIN | Finnish in Finland | HG00171 |
|  | GBR | British from England and Scotland | HG00096 |
|  | IBS | Iberian Populations in Spain | HG01500 |
|  | TSI | Toscani in Italia | NA20502 |
| **South Asian** | BEB | Bengali in Bangladesh | HG03006 |
|  | GIH | Gujerati Indians in Houston, Texas,<br>USA | NA20845 |
|  | ITU | Indian Telugu in the UK | HG03713 |
|  | PJL | Punjabi in Lahore, Pakistan | HG01583 |
|  | STU | Sri Lankan Tamil in the UK | HG03642 |

**Table S2.** Composition of the reference files used to test performance for new and sparsely-sampled populations.

| Reference<br>file | 1000G individuals<br>(no OCE) | OCE individuals |  |  |
| --- | --- | --- | --- | --- |
|  |  | PAS | BOU | PAH |
| Original | 2,478 | 0 | 0 | 0 |
| "5 OCE" | 2,478 | 5 | 0 | 0 |
| "10 OCE" | 2,478 | 5 | 5 | 0 |
| "15 OCE" | 2,478 | 5 | 5 | 5 |
| "20 OCE" | 2,478 | 5 | 10 | 5 |

**Table S3.** Real time for ADMIXTURE processing of a 30 sample file.

| # loci | Time (s) |  |  |
| --- | --- | --- | --- |
|  | Conversion step | Prediction step | Total |
| **1,000** | 2.64 | 1.26 | 3.90 |
| **2,000** | 4.24 | 1.93 | 6.17 |
| **5,000** | 9.09 | 4.60 | 13.69 |
| **10,000** | 17.24 | 10.13 | 27.37 |
| **20,000** | 33.46 | 20.16 | 53.62 |
| **50,000** | 81.99 | 47.99 | 129.98 |
| **100,000** | 163.40 | 154.99 | 318.39 |
| **128,097** | 212.39 | 220.08 | 432.47 |

### Figures

**Figure S1.**
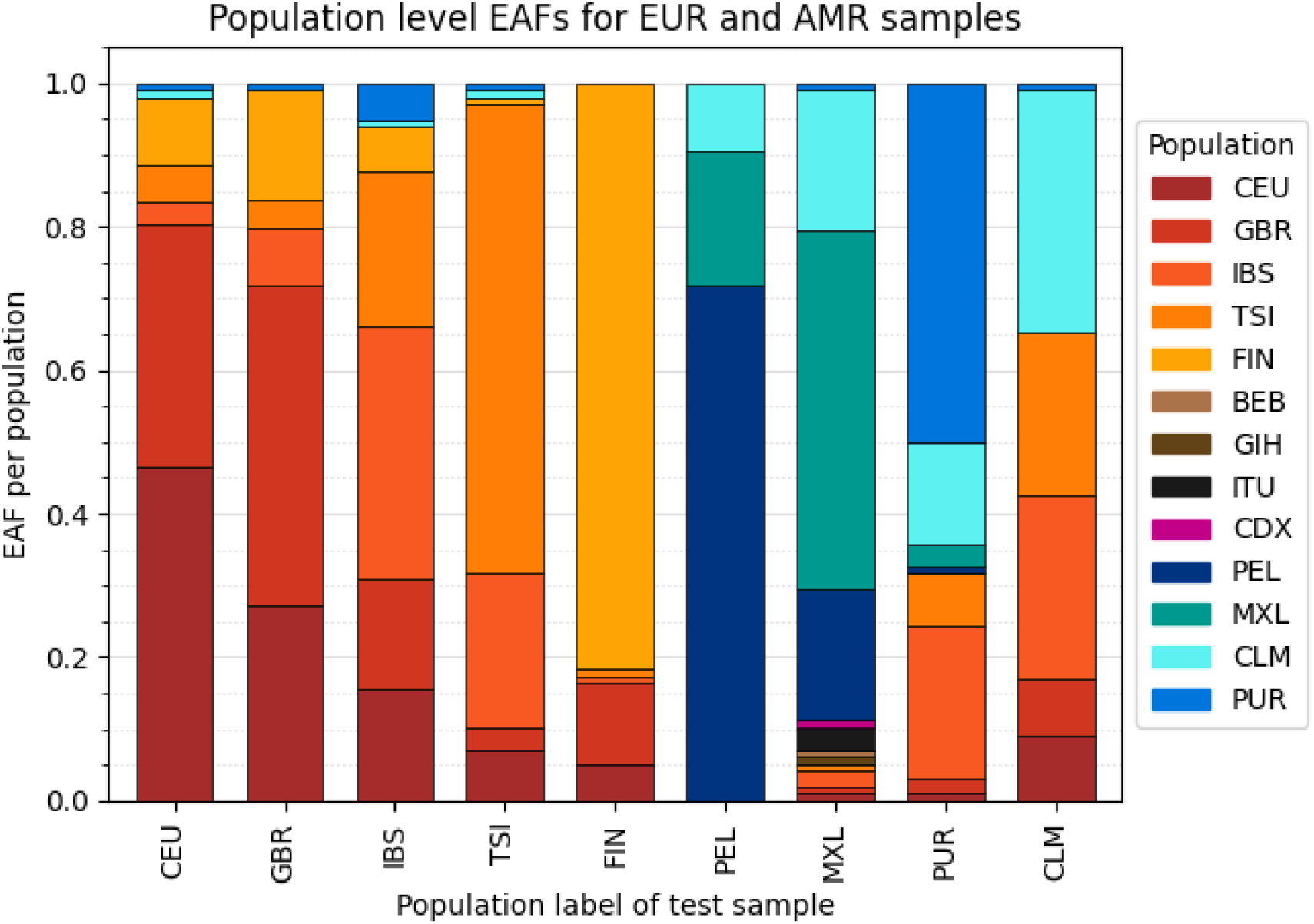
At 20,000 loci all samples apart from the CLM sample have the largest proportion of their ancestry assigned to the correct superpopulation (European: orange shades, American: blue shades). However, when viewed at the population level, the CLM sample is assigned a larger EAF to the CLM population than to any individual European population. We consider these proportions a reasonable estimate due to the historic movement of European peoples to this area [22,24]. The small proportion of ancestry assigned to American populations for the European test samples (CEU, GBR, IBS, TSI) likely reflects this admixed composition of American populations in the 1000 Genomes dataset used as a reference.

**Figure S2.**
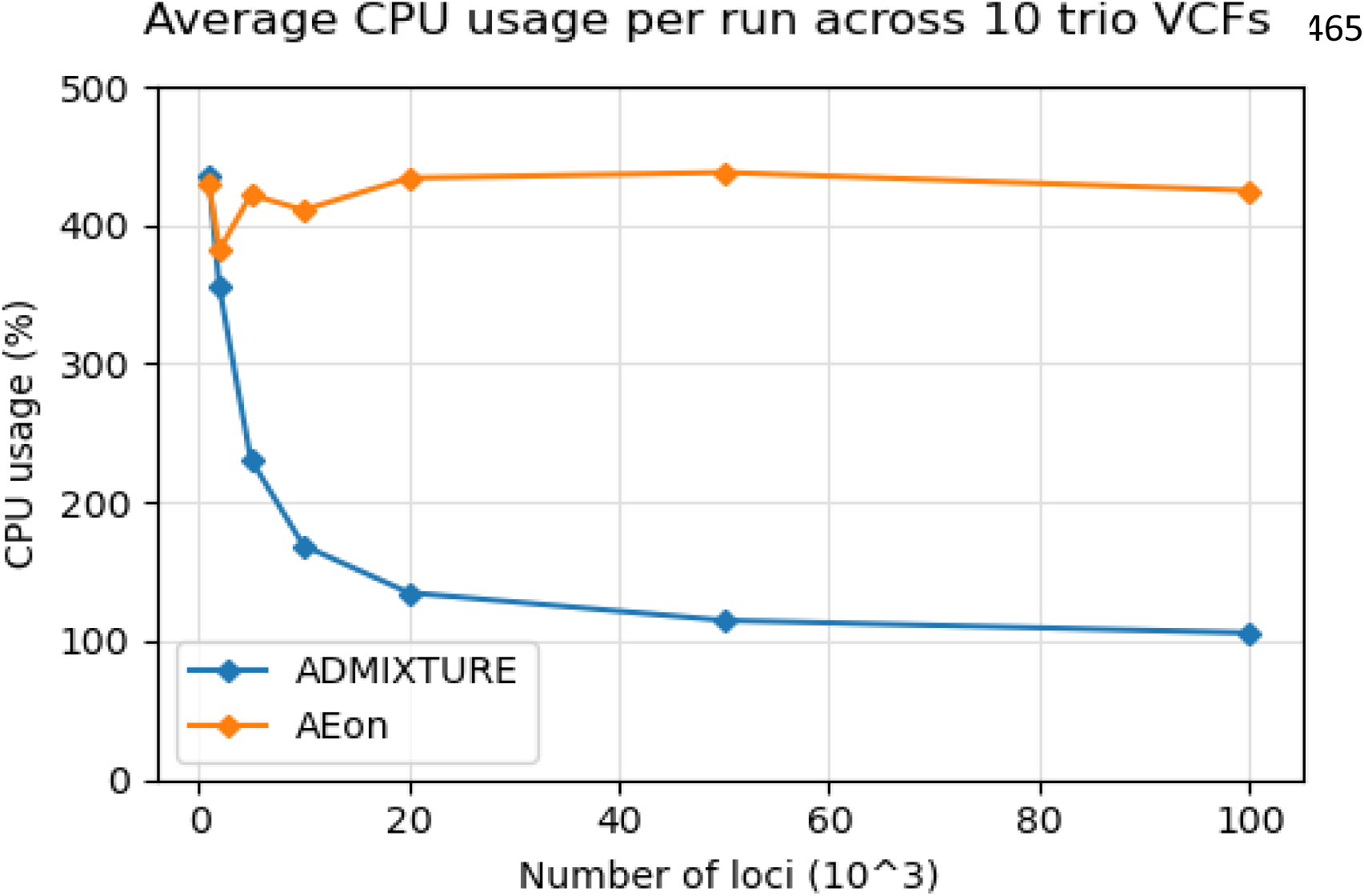
When run on files containing only three samples, ADMIXTURE was able to make use of the same amount of CPU as AEon with small number of loci.

**Figure S3.**
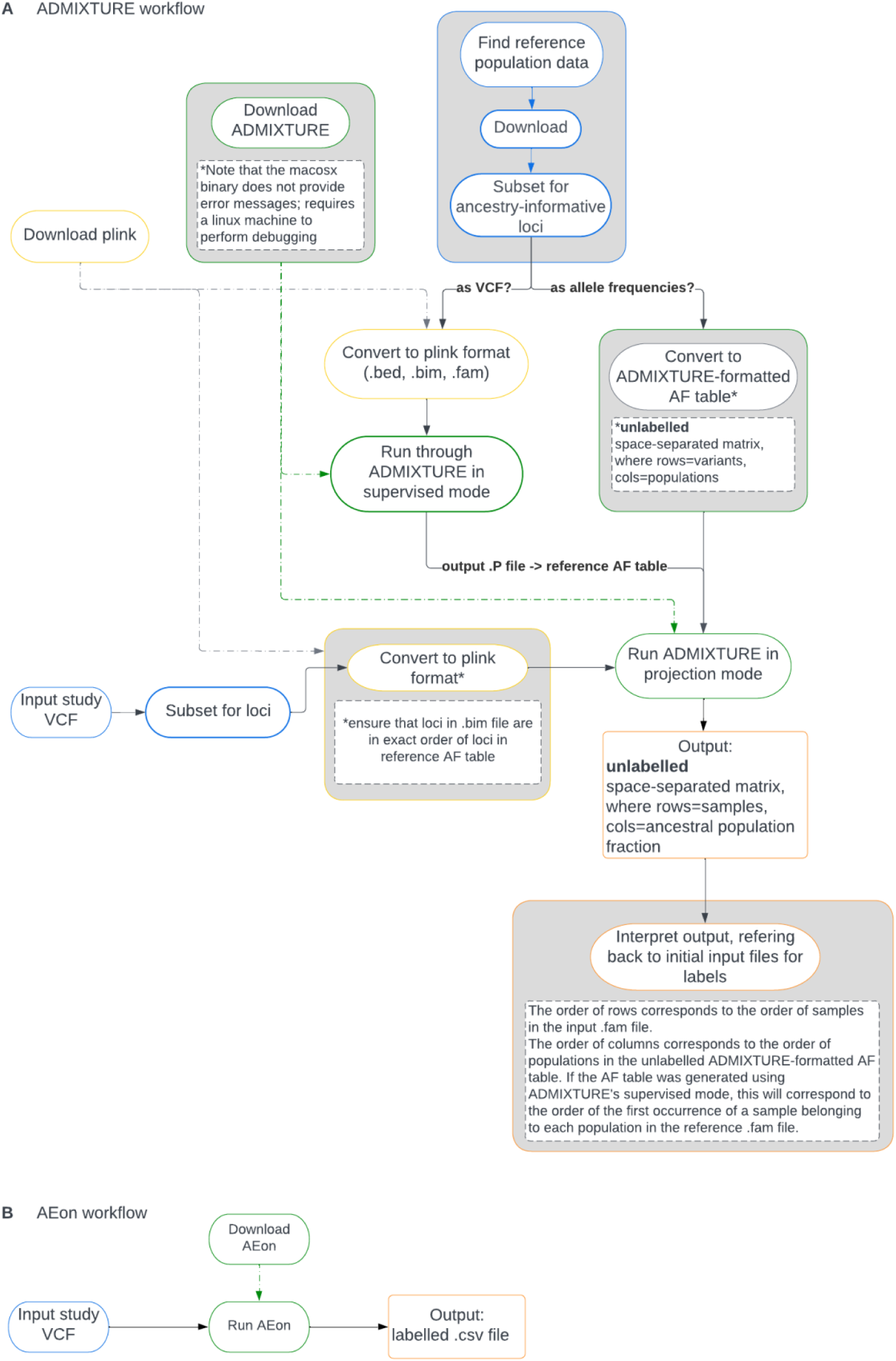
Flow chart demonstrating the process required to run ADMIXTURE compared with AEon.

**Figure S4.**
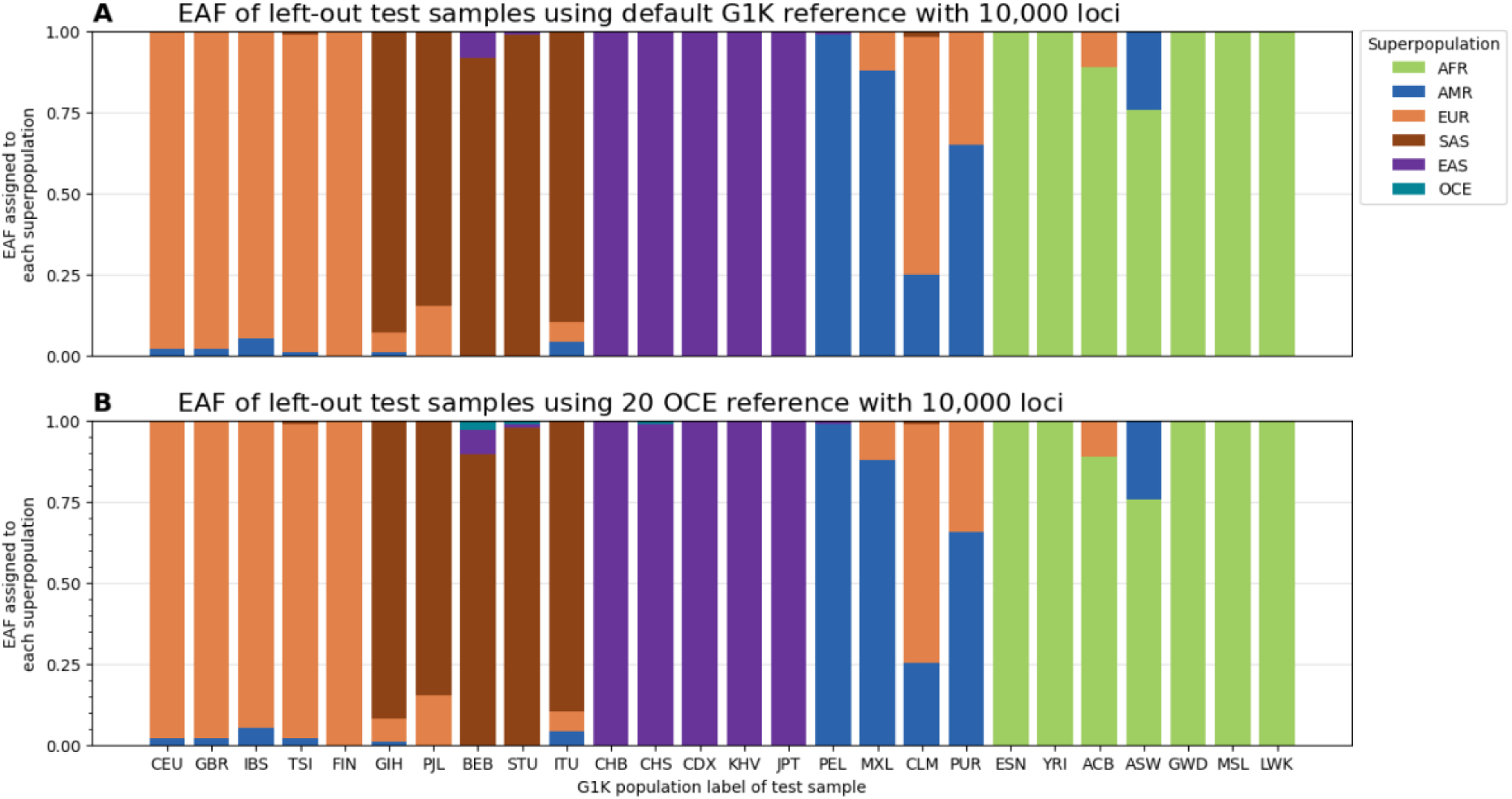
The addition of OCE samples to the reference file does not significantly impact the ancestry fractions assigned to samples from other populations.

### Additional files

**Additional file 1**

The pre-processing script used to convert BCFs into the input format required by ADMIXTURE is provided as *additional_file_1.sh*.

